# TopoDoE: A Design of Experiment strategy for selection and refinement in ensembles of executable Gene Regulatory Networks

**DOI:** 10.1101/2023.04.21.537619

**Authors:** Matteo Bouvier, Souad Zreika, Elodie Vallin, Camille Fourneaux, Sandrine Giraud-Gonin, Arnaud Bonnaffoux, Olivier Gandrillon

**Author notes:** Contributing authors.

## Abstract

**Background:** Inference of Gene Regulatory Networks (GRNs) is a difficult and long-standing question in Systems Biology. Numerous approaches have been proposed with the latest methods exploring the richness of single-cell data. One of the current difficulties lies in the fact that many methods of GRN inference do not result in one proposed GRN but in a collection of plausible networks that need to be further refined. In this work, we present a Design of Experiment strategy to use as a second stage after the inference process. It is specifically fitted for identifying the next most informative experiment to perform for deciding between multiple network topologies, in the case where proposed GRNs are executable models. This strategy first performs a topological analysis to reduce the number of perturbations that need to be tested, then predicts the outcome of the retained perturbations by simulation of the GRNs and finally compares predictions with novel experimental data.

**Results:** We apply this method to the results of our divide-and-conquer algorithm called WASABI, adapt its gene expression model to produce perturbations and compare our predictions with experimental results. We show that our networks were able to produce *in silico* predictions on the outcome of a gene knock-out, which were qualitatively validated for 48 out of 49 genes. Finally, we eliminate as many as two thirds of the candidate networks for which we could identify an incorrect topology, thus greatly improving the accuracy of our predictions.

**Conclusion:** These results both confirm the inference accuracy of WASABI and show how executable gene expression models can be leveraged to further refine the topology of inferred GRNs. We hope this strategy will help systems biologists further explore their data and encourage the development of more executable GRN models.

## Background

For the last 60 years, it has been commonly admitted that a precise knowledge of gene regulatory interactions is required to fully understand the processes of cell decision making (differentiation, proliferation or death) in response to a stimulus [1, 2]. Therefore, for the last three decades, the Systems Biology field has dedicated a great deal of effort to infer the structure of Gene Regulatory Networks (GRNs). Initial attempts suffered from the imprecision of bulk RNA-seq, in which the expression data from millions of cells was averaged, masking cellular heterogeneity and stochastic phenomena. Algorithms developed in the last ten years benefited from the development of single-cell RNA-seq technologies, which now allows to access mRNA distributions in more details and investigate causal dependencies between genes. Indeed, the single-cell resolution was shown to contain a much richer information that the mean value alone [3–5].

However, the precise identification of biological parameters of a GRN and the problem of distinguishing between multiple possible topologies remain to this day challenging problems. Attempts at solving those problems were for example made in the context of the DREAM challenges [6] where experimental design strategies were developed. The general goal of those strategies was to decide under which perturbation (gene knock-out (KO), knock-down (KD) or over-expression), at which time point(s) and through which kind of data (bulk or single-cell RNA-seq, proteomics, …) a process of interest should be observed to discard the largest amount of incorrect GRNs, therefore leading to a small number of most relevant GRNs (and ideally leaving only one). Those strategies must respond to the difficult question of maximizing the amount of newly acquired data while minimizing the costs (financial costs, time required), dealing with measurement uncertainties and accounting for the stochastic nature of gene expression.

Recently our team developed WASABI [7], a tool which allows to 1) infer GRNs from time-stamped scRNA-seq data and 2) simulate those GRNs. Simulations are made possible by a mechanistic model of gene expression, previously described in [8], where a stochastic process controls promoter activation and a set of ODEs determines RNA and protein synthesis, resulting in a Piecewise Deterministic Markov Process (PDMP). The algorithm works by iteratively building and simulating ensembles of candidate GRNs from which the best performing are selected.

This GRN inference algorithm was applied to a dataset of single-cell RTqPCR data obtained on differentiating chicken erythrocytic cells. As expected, it did not produce a single GRN but rather a collection of 364 candidate GRNs equally well suited for reproducing experimental data when simulated. It is therefore the ideal playground for the development of a Design of Experiment strategy able to efficiently reduce the number of candidate GRNs previously generated by a GRN inference algorithm.

To do this, we introduce TopoDoE, an iterative method for the *in silico* identification of the most informative perturbation – that is eliminating as many incorrect candidate GRNs as possible from the data gathered in one experiment. That method is a 4 step process in which :

1. a topological analysis is performed on the set of candidate GRNs to identify the most promising gene targets. This is essential to avoid the heavily time-consuming simulation of all possible gene perturbations.
2. *in silico* perturbation and simulation of the identified gene targets and ranking of those perturbations to identify the most informative one.
3. *in vitro* execution of the selected perturbation and scRNA-seq data acquisition.
4. selection of the subset of candidate GRNs which accurately predicted the novel experimental data.

This strategy led to the identification of the *FNIP1* gene as a promising target, that was knocked-out in chicken erythrocytic progenitor cells. The *in silico* predictions of *FNIP1* KO were verified for 48 out of 49 genes in our GRNs. The DoE strategy helped reduce the 364 candidates into 133 most relevant ones. The merging of those 133 GRNs led to one GRN with a much improved goodness of fit to experimental data than any other candidate.

## Results

### Initial setting

WASABI has previously been applied to the inference of the GRN governing the differentiation process of avian erythrocyte progenitor cells (T2ECs) into mature erythrocytes [7]. It generated 364 candidate GRNs, all made of the same 49 genes (**S1 Table**) and of a unique stimulus mimicking the change of culture medium which triggers the differentiation process [9]. As shown in **S1 Fig**, all 364 GRNs shared an overall close but always different topology. When comparing all GRNs two-by-two, we found on average a low number of different interaction values between pairs of genes: only 7.72 different values out of the 160 total existing interactions. Fig 1.A shows the graph of interactions of one such candidate GRN.

**Fig. 1.**
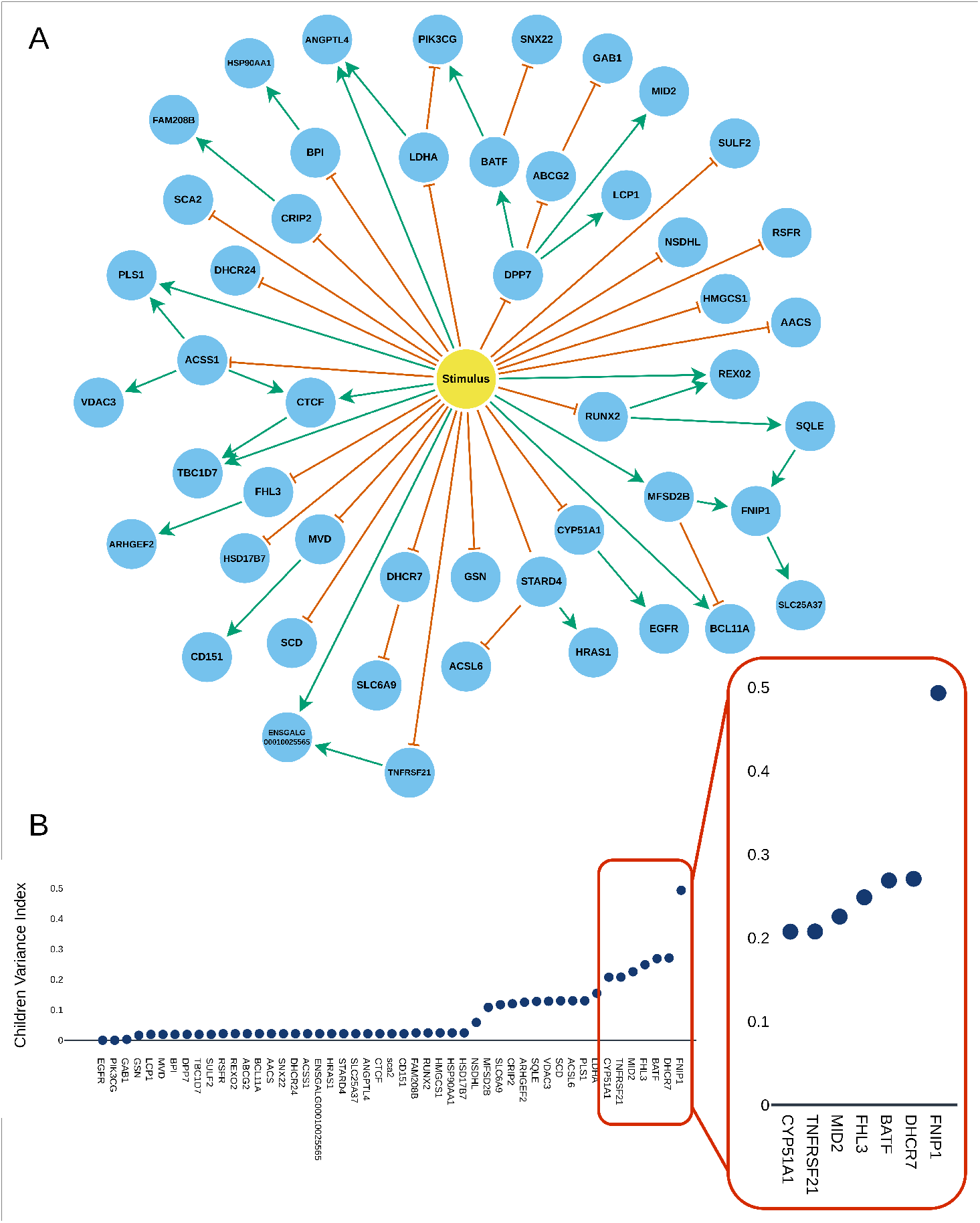
Topological features of the 364 candidate GRNs inferred by WASABI. **A** Graph of one of the 364 candidate GRNs. Genes are shown as blue nodes and the stimulus as a yellow node. Green edges represent positive regulations (*θ >* 0) from a gene source to a gene target while orange edges represent negative regulations (*θ <* 0). **B** Descendants Variance Index (DVI) per gene. This index gives the variance of interactions between a gene and the genes it regulates, found in all 364 GRNs. A high value indicates that a gene has highly varying interactions among all of the candidate GRNs.

GRNs generated by WASABI were defined by a mechanistic model of gene expression based on coupled Piecewise-Deterministic Markov Processes (PDMPs) governing how the mRNA and Protein quantities change over time. In this model, the gene promoter activation (i.e. gene bursting frequency) is function of the expression level of all other genes. Gene *A* is said to regulate gene *B* when the interaction value *θ*_*A,B*_ is not null.

Because this model was executable, it allowed us to simulate the behavior of GRNs over some period of time by solving the underlying PDMPs. The result of a GRN simulation was a collection of matrices of cells *×* genes values of mRNA counts, one for each time point. When simulated, all 364 candidate GRNs produced similar count matrices: distances between simulated and experimental data were indeed all close (see **S2 Fig**), with distance variations explained purely by the randomness of the simulations and no GRN performing significantly better than others. Here, distances were computed using the Kantorovich distance [10] taken on marginals (i.e. computed one gene at a time). Our next objective was thus to identify a perturbation which would produce different GRN responses in the form of diverse count matrices.

#### Step 1: Topological analysis

Depending on the number of genes in the GRNs of interest, simulation of all possible perturbations on all genes might be very time consuming or even completely unfeasible. We thus sought to develop a preliminary step to our strategy, based on the topological analysis on the set of candidate GRNs, that would allow us to identify genes having the highest chance of producing informative perturbations. This analysis was motivated by the fact that, while some gene-to-gene (or stimulus-to-gene) interactions appeared in all GRNs, others were present in very few of them (for example, the regulation of *FNIP1* by *GAB1* only appears in one of the 364 candidates; see **S3 Fig**). In particular, the gene *FNIP1* had many possible regulator genes : interactions with 47 out of the 49 genes could be found in the set of candidates, but only in at most 9 GRNs at a time (except for the regulation by *MFSD2B* which was found in all GRNs). This configuration is intuitively promising since the GRNs would produce many different regulatory dynamics upon perturbation of a single target gene, because of their many distinct gene-to-gene interactions.

To identify the genes with the most variable interactions with its descendants (the downstream genes it regulates), we proposed the **Descendants Variance Index (DVI)**. Briefly, this index considers one gene at a time and measures how much interactions between that gene and its descendants **qualitatively** change in the whole set of candidate GRNs (change from activation to inhibition or to no interaction at all). High values for a gene on the DVI indicate that many different types of regulations can be found in the GRNs while low values show that most GRNs have the same regulations. Here, we focused only on downstream genes since we will only consider KO experiments in later steps, thus only affecting the expression of genes regulated by the KO target, as it is the case for most kinds of perturbations.

DVI values where highest for genes *FNIP1* (DVI=0.4934), *DHCR7* (DVI=0.2707), *BATF* (DVI=0.2687), *FHL3* (DVI=0.2487) and *MID2* (DVI=0.2255), as shown in Fig 1.B. Those 5 genes were thus selected for studying the effect of their *in silico* perturbation.

#### Step 2: *In silico* perturbations

We simulated the 364 candidate GRNs after KO of each of the genes identified in the previous analysis. Simulated data had to be compared to reference unperturbed data to obtain predictions on which genes would display significant expression variations upon perturbation. Finally, a measure of entropy was used to select the most informative perturbation.

Before any simulation could be run however, constant hyper-parameters of our model needed to be chosen so as to obtain balanced simulations. The goal here was to set the initial state of the simulations so that they would reproduce experimental data from unperturbed cells in a stable way in the absence of any perturbation. This was an essential preliminary step for correct interpretation of the results since incorrect simulation balancing would introduce simulation biases, either spontaneously drifting away from the initial state or remaining stuck on it and thus masking the effect of a perturbation. We devised an initialisation method based on a modified simulated annealing algorithm which simulated gene expression values for 40 hours with different hyper-parameters’ values to identify optimal combinations.

Then, *in-silico* data was obtained for all 6 conditions (reference data with no perturbation and each of the 5 selected genes knocked-out independently). For each, all 364 candidate GRNs were simulated for 100 hours, so that they had enough time to reach a new stable state after perturbation. Data was recorded at the end of the 100 hours to obtain mRNA count matrices of 200 cells each.

Counts of mRNA molecules after each perturbation were compared to the reference dataset using one-sided t-tests for both ‘greater’ and ‘less’ mean value hypothesis. The effects on downstream genes are shown in Table 1, where the number of GRNs for which we measured significant expression variation is reported (p-value *<* 0.01). Rows colored in red indicate uninformative effects of a KO. Indeed, when the effect of a perturbation was the same in all of the 364 candidate GRNs or when no GRN responded to that perturbation, it provided no valuable information to discriminate GRNs. To assess the total amount of information given by a KO experiment, we measured for each gene the entropy on the proportions of GRNs with variations in expression levels. Measured entropy values are given in Table 2. The KO of *FNIP1* had the largest entropy (i.e. carried the most information) with a value of 1.0466. This perturbation was thus selected for the next steps.

**Table 1.** Effects of *in-silico* perturbations. The effects of single gene KO’s on downstream genes were tested using one-sided t-tests for ‘less’ expression in the KO condition (**down-expression** in the ‘Effect’ column, p-value *<* 0.01) and for ‘greater’ expression in the KO condition (**up-expression** in the ‘Effect’ column, p-value *<* 0.01). The number of GRNs in which the gene showed an expression level variation is stored in the column ‘GRNs’. Values in parenthesis indicate the number of GRNs which had a non-null interaction between the knocked-out gene and the downstream gene, which is the maximum expected number of GRNs showing an expression level variation. Asterisks indicate uninformative expression variations on downstream genes.

| Knock-Out target | Gene | Effect | GRNs (with interaction) |
| --- | --- | --- | --- |
| FNIP1 | SNX22 | down-regulation | 88 (88) |
|  | SLC25A37 | down-regulation | 45 (45) |
|  | ENSGALG00010025565 | up-regulation | 102 (102) |
| DHCR7 | FNIP1 | down-regulation | 6 (8) |
|  | SLC6A9 | up-regulation | 1 (192) |
| BATF | FNIP1 | down-regulation | 7 (7) |
|  | PIK3CG | down-regulation | 364* (364) |
|  | SNX22 | up-regulation | 178 (178) |
| FHL3 | ARHGEF2 | down-regulation | 364* (364) |
|  | FNIP1 | down-regulation | 8 (8) |
| | SLC6A9 | ∅ | 0* (127) |
| MID2 | FNIP1 | down-regulation | 7 (8) |
| | SLC6A9 | ∅ | 0* (45) |
|  | ENSGALG00010025565 | down-regulation | 39 (39) |

**Table 2.** Measured values of entropy after single gene KO.

|  | *FNIP1* | *BATF* | *MID2* | *DHCR7* | *FHL3* |
| --- | --- | --- | --- | --- | --- |
| entropy | 1.0466 | 0.6933 | 0.4265 | 0.4101 | 0.1038 |

### Predictions of gene expression variation

Since we had selected *FNIP1* as target for a KO experiment, we sought to obtain *insilico* predictions of the expected gene expression variations. To that end, the mRNA counts from all GRNs simulated under *FNIP1* ‘s KO were pooled together and compared to the reference condition to produce average predictions. As shown in Fig 2 and as expected from the results in Table 1, genes *SNX22* and *SLC25A37* had overall decreased mRNA counts in the KO condition and *ENSGALG00010025565* had increased mRNA counts (one-sided t-tests with p-value *<* 0.01). Also as expected, none of the 45 other genes had a significant expression variation (see **S4 Fig**).

**Fig. 2.**
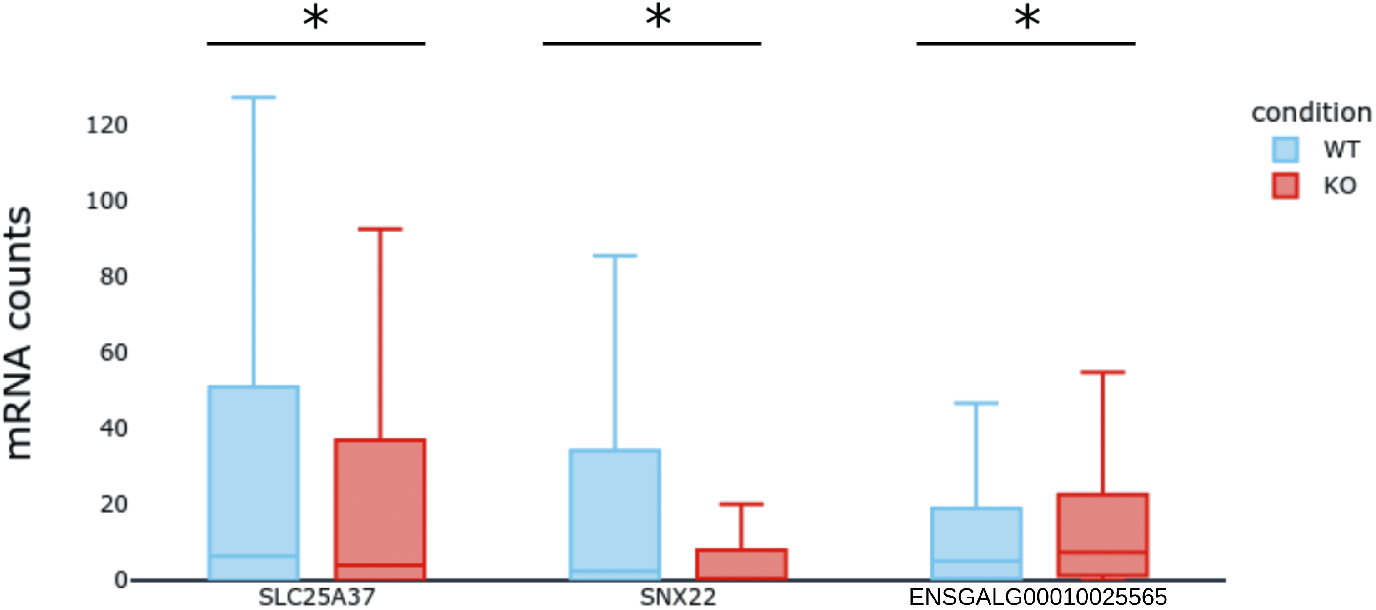
Simulations of FNIP1’s Knock-Out. Overall predictions on the gene expression levels after the *in silico* KO of FNIP1. Box plots summarize the mRNA counts obtained from the simulation of the 364 candidate GRNs in the Wild Type (WT, in blue) and in the knock-out (KO, in red) conditions, after 100 hours of simulation. Shown are the three genes with significant (p *<* 0.01) expression variation.

#### Step 3: *In vitro* perturbations

To test experimentally our *in-silico* predictions, we devised a dedicated strategy to obtain single cell transcriptomics data on cells that had been validated for the KO of *FNIP1* (Fig 3.A). One of the main challenges we had to face was that the T2ECs being primary cells, they have a finite lifetime of 30 days [9] during which the cells had to be transfected, cloned, amplified, molecularly validated, and seeded in 96 wells plate for subsequent scRTqPCR analysis.

**Fig. 3.**
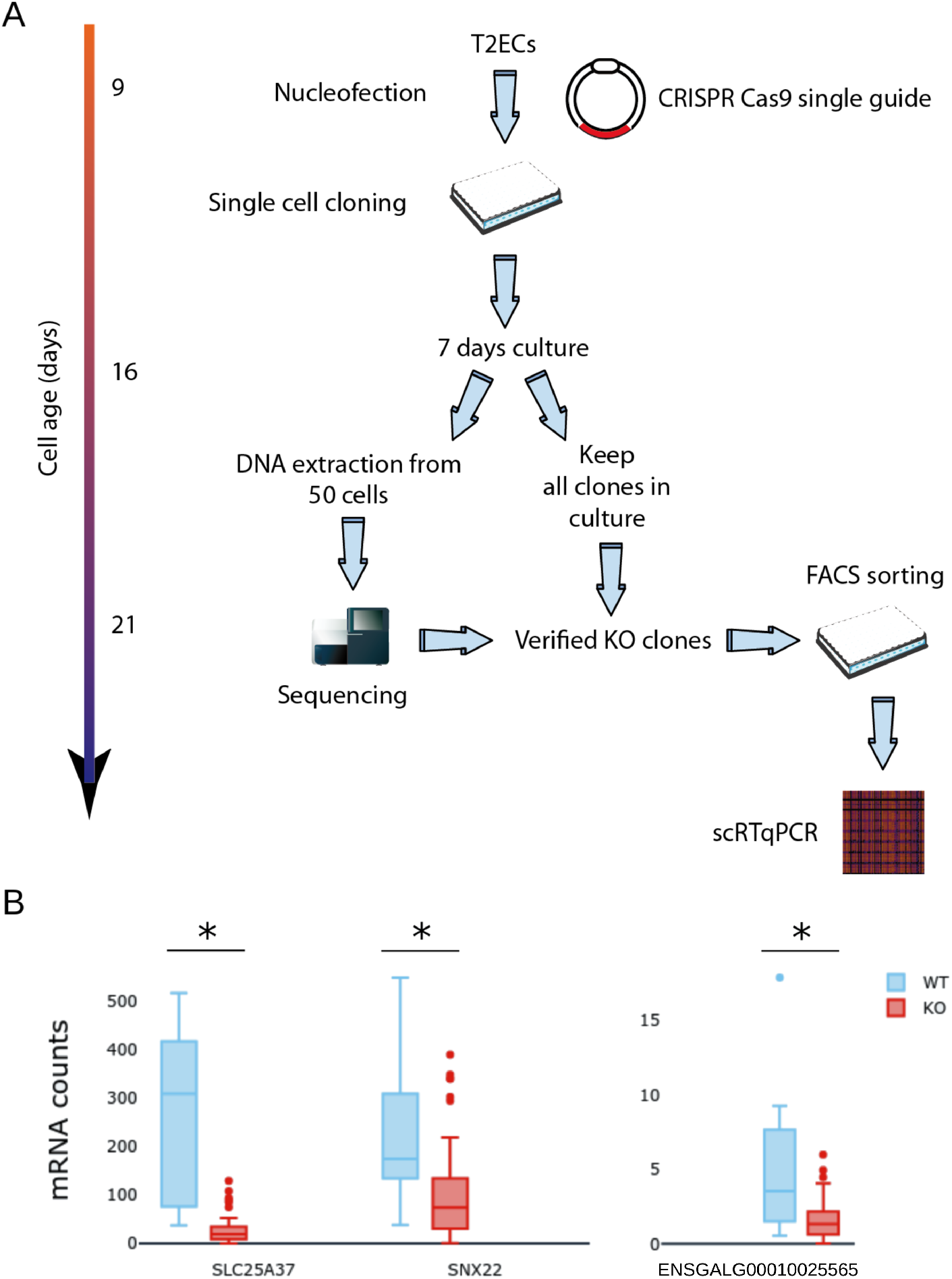
Generation and single-cell analysis of *FNIP1* KO cells. **A** The experimental strategy used for generating scRTqPCR data on validated *FNIP1* KO cells. **B** Single cell counts for wild type cells (in blue) and *FNIP1* Knock-Out cells (in red).

Following *FNIP1* ‘s KO, we acquired single cell transcriptomics data for 61 KO cells and for 12 cells transfected with an empty plasmid which we used as control (Fig 3.B). We recovered expression data for 45 of the 49 genes in the GRNs. Genes *ABCG2, LDHA* and *GAB1* displayed poor quality data and were removed from the dataset.

Using one-sided t-tests, we found that the expression of genes *SNX22* and *SLC25A37* dropped significantly in the KO condition when compared to the control, which matched our predictions (see **S2 Table** and **S5 Fig**). Surprisingly however, the expression also dropped for genes *ENSGALG00010025565* and *SLC6A9*. This indicated a flaw in WASABI’s inference method where it overestimated the basal expression level or the auto-activation strength for gene. For example, *ENSGALG00010025565* ‘s expression level was often supported by it’s own expression alone, as much as to not be regulated by any other gene in some candidate GRNs.

Importantly however, as previously predicted, no significant expression level variation was measured for the 45 other genes. Altogether, these results allowed us to confirm that WASABI was able to infer mostly correct GRNs with a remarkable accuracy. Indeed, the probability of making at most 2 errors on the qualitative responses of the 49 studied genes was only *P* (*E* ≤ 2) = 2.0071 · 10^−20^ with *E* the number of errors and with a probability of 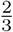 of making an error for each gene.

#### Step 4: GRN selection and refinement

From the previous step, we had identified a KO target, generated predictions on the gene expression variations after perturbation and verified most of our predictions from experimental KO data. The last step of our strategy was to rule out incorrect GRN candidates. From the novel information gathered in the experiment, we were also able to build a GRN more accurately reproducing the data.

GRNs were selected by retaining only those with topologies coherent with the obtained experimental results. 45 GRNs with *FNIP1* positively regulating *SNX22* matched the decreased expression of *SNX22* and 88 GRNs with *FNIP1* positively regulating *SLC25A37* matched the expression drop of that gene. We decided to select those 133 GRNs among the 364 candidates and rule out the 231 others.

Interestingly, no GRN had *FNIP1* regulating both *SNX22* and *SLC25A37* simultaneously. This revealed a limitation in WASABI’s exploration of possible GRN topologies which was caused by the limited computational resources available at the time WASABI was developed. This limitation prevented WASABI from exploring more complex topologies in which *FNIP1* regulated more that one gene at once. To overcome this limitation, the 133 selected GRNs were merged into a single GRN by computing the average of interaction values for each stimulus-to-gene or gene-to-gene regulation.

To measure the performance of this new GRN, we computed the distance of simulated to experimental KO data for all 364 candidate GRNs. The distribution of such distances is shown as the blue histogram in Fig 4. Similarly, we computed the distance of simulated data obtained with the merged GRN to the experimental data (green dotted line). This distance was much lower than those obtained with any other GRN, demonstrated the improved goodness of fit of the constructed GRN.

**Fig. 4.**
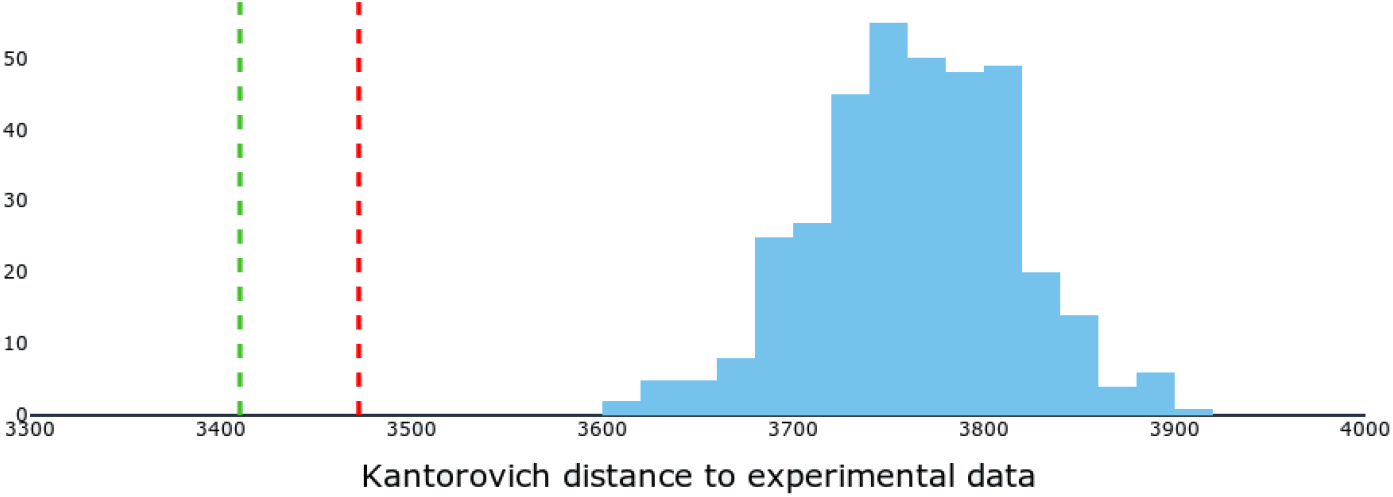
Simulations of *FNIP1* ‘s Knock-Out. Evaluation of the GRNs’ goodness of fit to experimental data. Kantorovich distances between experimental and simulated data were computed for the 364 candidate GRNs. The blue histogram shows the distribution of those distances. In red is the distance obtained from the simulation of the 364 GRNs merged into one. In green is the distance obtained after merging into one the 133 selected GRNs.

Finally, to verify the relevance of our selection, we merged all 364 candidate GRNs in the same way as before and again computed its distance to the experimental data (red dotted line in Fig 4). Again, we obtained a better performance than any of the 364 GRNs, which could be explained by the fact that, by merging all of the GRNs, we recovered the simultaneous regulation of *SNX22* and *SLC25A37* by *FNIP1*. However, that merged GRN did not perform as well as the one obtained by merging the 133 selected GRNs, confirming the relevance of our selection.

## Discussion

We have presented TopoDoE, a DoE strategy that was designed for selecting the most informative experiment to perform to significantly reduce the number of previously inferred GRNs. When applied as a follow-up step to WASABI’s GRN inference algorithm, the presented strategy of network selection allowed to first identify and remove incorrect GRN topologies and then to recover a new GRN better fitting experimental data than any other candidate.

### Validation of the inference algorithm

Initial simulations of GRNs inferred by WASABI showed they all fitted equally well the experimental data used in the inference step. This motivated the generation of new experimental data able to distinguish between the candidate networks. The simulation and then the experimental completion of *FNIP1* ‘s KO further showed that the 364 candidate GRNs proposed by WASABI closely matched the “true” GRN. Indeed, among the 49 studied genes, the expression level variation after *FNIP1* ‘s KO was incorrectly predicted for only 1 gene (*ENSGALG00010025565*). Most importantly, the sole up-regulations of *SNX22* and *SLC25A37* were anticipated, indicating that no other interactions with *FNIP1* as regulator were missed during the inference step. The probability of generating GRNs which would produce such accurate predictions purely by chance was extremely low. This finding provides us confidence on the quality of both WASABI’s algorithm and the inferred GRNs.

### Identification of WASABI’s limitations

One interesting finding was the ability of TopoDoE to also identify limitations in the GRN inference algorithm. A closer inspection of the candidate GRNs indeed revealed two main issues in the initial implementation of WASABI : (i) the exploration of possible GRN topologies was incomplete and (ii) selected topologies has a bias towards strong auto-activation regulations.

When WASABI was run on a super computer, it required one entire CPU node per tested topology for its simulation. Combined with the simulation slowness, this lead to high needs for computation resources which in turn meant that too few GRN topologies could be explored per iteration of the algorithm. WASABI thus could not explore more complex alternatives in which *FNIP1* would regulate more than one gene at a time. This is a common issue in the general scope of Machine Learning in which **exploration** – testing different solutions – and **exploitation** – evaluating a particular solution’s relevance – need to be correctly balanced (see for example [11] or [12]).

Also, WASABI allowed some genes to have high basal expression level supported by strong auto-activations. In some cases, genes such as *ENSGALG00010025565* were in fact only regulated in that way and were thus completely disconnected from the rest of the GRN. This evidently prevented the prediction of the positive regulation of *ENSGALG00010025565* by *FNIP1* as shown in the experimental data. This behavior can easily be corrected by adding a penalisation term to auto-activations in the future use of WASABI.

### Quality of the predictions

Even though *SNX22* ‘s and *SLC25A37* ‘s variation of expression levels after *FNIP1* ‘s KO were statistically significant in both simulation and experimental data, it must be noted that the predictions were only qualitatively accurate, but not quantitatively. Indeed, expression levels for both genes were very low in simulated data as compared with experimental data of *FNIP1* Knock-Out. This observation can be explained by the variability in mRNA counts between experiments. Indeed, mRNA counts did not always perfectly coincide between the training data (used to infer the GRNs) and the experimental data obtained after *FNIP1* ‘s KO. Also, the small number of cells in the experimental data (12 wild type and 61 knocked-out cells) could have induced a bias.

### GRN selection and refinement

In the final step of our strategy, we selected the candidate GRNs which qualitatively predicted the expression variation for at least one gene, after *FNIP1* ‘s KO. This resulted in a total of 133 selected GRNs, thus eliminating two thirds of the candidates.

Even though the “true GRN” was not in the initial set of candidates, it was possible to recover it at least partially by merging promising GRNs together. Here, when merging the 133 selected candidates, we built a new GRN which performed significantly better than all other candidates in reproducing the reference data.

### Expanding to other perturbations

In this work, we only worked with KO perturbations since we had mastered the CRISPR-cas9 system in T2ECs, which allowed to perturb all genes downstream of the target. However, TopoDoE could easily be expanded to other types of perturbations such as knock-downs (where short interfering RNA fragments inhibit the translation of specific mRNAs [13, 14]) or over-expressions (obtained by introducing a dedicated plasmid in a cell [13]). In some cases, these approaches would be easier to apply than full knock-outs or allow to perturb genes that cannot be knocked-out because they are essential for the cell’s viability. Knock-downs and over-expressions can also be easily introduced in our expression model by increasing by some factor the d0 and s0 parameters respectively. Here we would focus on d0 and s0 to stay close to the biological processes since knock-downs increase the rate of mRNA degradation while over-expressions increase the amount of mRNA molecules synthesized.

Interestingly, we found that the variability in interaction values among the candidate GRNs was maximal not when considering interactions between FNIP1 and the genes it regulated, but when considering interactions with the genes that regulated FNIP1 (see **S6 Fig**). It is however difficult to devise a perturbation targeting at once all of the interactions between a gene and those upstream of it, thus producing the maximum amount of different GRN responses.

One possibility might be to use a reversion experiment, in which the differentiation stimulus is interrupted early. Depending on the exact regulation dynamics, the expression level of *FNIP1* could change greatly. From previous work [15], we know that T2ECs definitely commit to the differentiation after a precise amount of time after stimulation, but remain able to fully revert to the progenitor state before that. The exact time at which the commitment happens heavily depends on the GRN’s topology and would thus allow to discriminate between different candidate topologies.

### Iterating the DoE strategy

Our strategy can be repeated iteratively to further decrease the number of candidates: the set of selected GRNs could now be used as input for the topological analysis step. With a sustained rate of GRN selection of 50 to 70%, about 5 cycles of our strategy would be needed to reduce the original set of 364 GRNs to only 10 candidates. This would dramatically improve our confidence on the topology of the true GRN and thus greatly improve the precision of our predictions.

Finally, one should note that this strategy is not restricted to WASABI-generated GRNs, but is applicable whenever an ensemble of executable models of GRNs can be obtained.

### Applicability

As discussed in this work, TopoDoE heavily relies on the simulation of sets of candidate GRNs. To apply it in other settings, it is thus necessary to have produced ensembles of executable GRN models. Although few such models currently exist, several executable models have been published in the last years. Many of such models come from the field of Boolean networks in which GRNs are executable by nature [16, 17]. Additionally, some models use an internal model of reactions to simulate single cell data accurately [18] as it is done in WASABI.

Finally, ensemble inference algorithms are still very uncommon to our knowledge, with notable exceptions such as [19] and [20]. This shows that the inference of ensembles of executable GRNs is a very valuable characteristic of WASABI. Our DoE strategy still remains applicable by combining inference and simulation methods cited above and the growing interest for executable ensembles of GRNs makes us believe that more such algorithms will come in the next years.

### Conclusions

Inference of GRNs has been at the heart of the systems biology field for decades, with numerous algorithms having been proposed. Their success has however been only partial because of the extreme complexity of the problem. In recent years, important progress has been made by :

- exploiting the richness of single-cell transcriptomics data
- introducing executable models of gene expression
- inferring ensembles of GRN topologies at once

Our strategy was specifically designed for such settings, where the goal – after some GRN inference process has produced and ensemble of executable candidate GRNs – is to identify where information is lacking for a more precise identification of the true GRN topology.

TopoDoE is divised into 4 simple steps, each aimed at 1) reducing the complexity of identifying informative perturbations, 2) generating predictions on the effects of a perturbation and selecting the best perturbation, 3) collection of experimental data after perturbation and finally 4) comparison with predictions for GRN selection.

In this work, we limited our demonstration to single gene knock-outs but our strategy can easily be expanded to any kind of perturbation that can be simulated with the leveraged gene expression model. This, together with its iterative nature, makes us believe that our strategy has the potential to be used by many biologists wishing to refine their knowledge on the GRN they are studying.

It is also important to note that our results confirmed the remarkable efficiency of WASABI. Average predictions from the full ensemble of candidate GRNs proved to be correct for 48 out of the 49 genes in the networks, after the gene knock-out. With corrections made to the identified limitations of the algorithm, this gives us great confidence in WASABI and TopoDoE’s ability to build high quality GRNs, providing us with a tool for efficiently exploring and understanding complex cellular processes and diseases.

## Methods

### Average number of differences between GRNs

To measure the number of differences between candidate GRNs topologies, we considered GRNs as directed graphs where nodes were the genes and edges were the *θ* interaction values. We computed for each pair of GRNs *a* and *b* the number *d*_*a,b*_ of different *θ* values between all pairs of genes *i* and *j* :

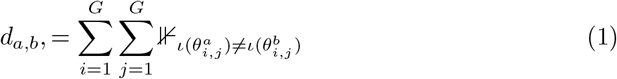

with

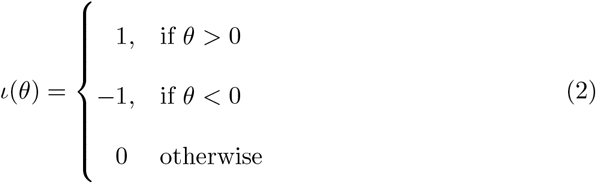

where *G* is the number of genes in the GRNs and 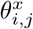 is the interaction value between genes *i* and *j* in GRN *x*. All *d*_*a,b*_ values were finally averaged to obtain the mean number of pairwise differences.

### Mechanistic model of gene expression

Simulations were run using a mechanistic model of gene expression described in [8] and based on the two-state model. Briefly, a gene *i* is described by a promoter *E*_*i*_ which can be in states *on* or *off* and randomly switches between those states at rates *k*_*on,i*_ and *k*_*off,i*_ respectively. When the promoter is active (*on* state), mRNA molecules (*M*_*i*_) are synthesized at rate *s*_0_, *i*. At any time, proteins (*P*_*i*_) are produced at rate *s*_1_, *i* from mRNA molecules, mRNAs are degraded at rate *d*_0_, *i* and proteins are degraded at rate *d*_1_, *i*. The following equations summarize the model :

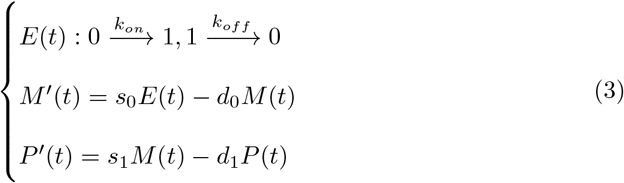

Interactions between stimuli and genes in a GRN are encoded by interaction parameters *θ* and by letting rates *k*_*on,i*_ and *k*_*off,i*_ be functions of protein *P* = (*P*_1_, …, *P*_*G*_) and stimuli *Q* = (*Q*_1_, …, *Q*_*S*_) levels as described in equation 4 (see Fig 5).

**Fig. 5.**
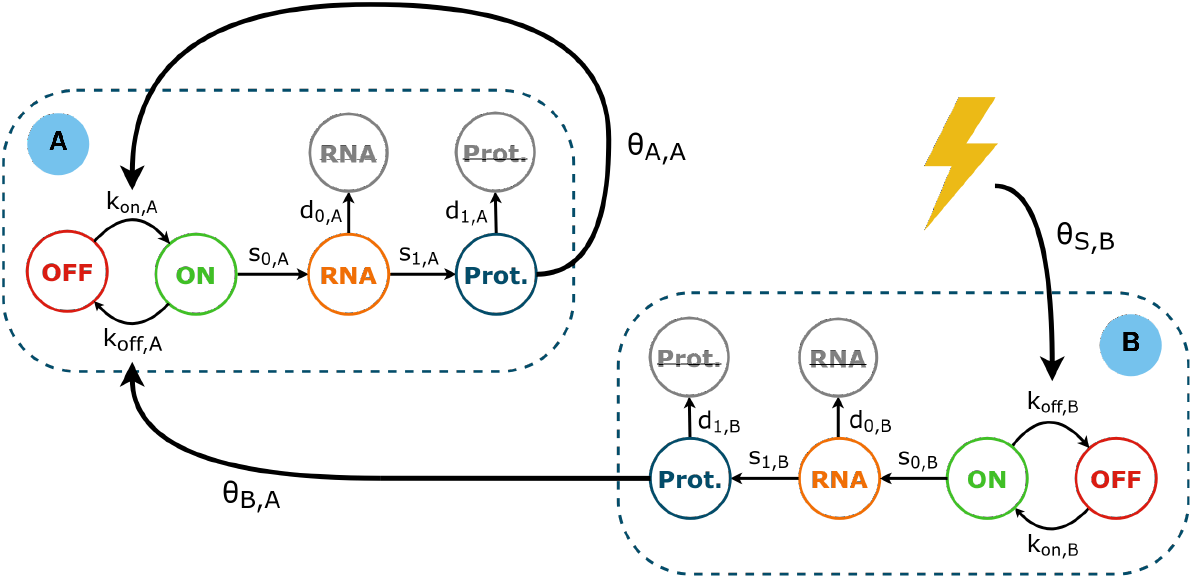
Executable GRN model. An example of a model of 2 genes **A** and **B** with a stimulus **S** represented by a yellow thunderbolt. Gene B regulates genes A, shown here by a non-null *θ*_*B,A*_ value, and is itself under the regulation of the stimulus. Gene A has an auto-activation loop : *θ*_*A,A*_ defines a self regulation.

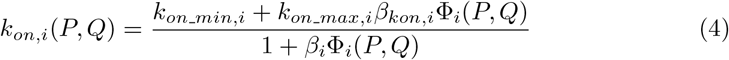

With

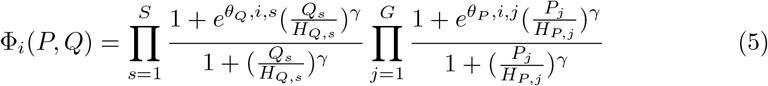

In equation 5, *H*_*Q,s*_ and *H*_*P,j*_ are interaction thresholds for stimuli and proteins respectively. Similarly, the interaction function for the rate *k*_*off*_ is given by equation 4 when replacing occurrences of index _*kon*_ by _*koff*_. Equation 5 is a modified version of those introduced in [8] and [7] to account for multiple stimuli and with the added stimulus threshold parameter *H*_*Q,s*_. The Hill exponent parameter *γ* is set to 4 in all cases.

For each gene, parameters *β*_*kon,i*_ and *β*_*koff,i*_ modify the gene’s basal expression level to account for the constant influence of genes outside of the modelled GRN. *β* values are estimated from experimental single-cell mRNA distributions but their correct identification is challenging and the algorithm used to that end is described in section Simulation balancing.

### Knock-Out perturbation implementation

Gene KOs were implemented in the simulation model by setting all *θ* interaction values between the perturbed gene and its neighbors to 0. *β*_*Kon*_ and *β*_*Koff*_ values were also set to 0 for that gene. During simulation, the probability of promoter activation (E being in state *on*) was forced to 0 so that the gene would not even be transcribed at the basal level.

During balancing and simulation, reference mRNA and protein counts were systematically set to 0 for the knocked-out gene so that the simulation would start completely devoid of such molecules.

### Variance Indices

Per-gene variances in the gene-to-gene and stimulus-to-gene interactions in a set of GRNs (all sharing the same set of genes and stimuli) were computed using a variance index. First, all interaction values were categorized into activation (if the interaction value was greater than 0), inhibition (if the interaction value was lesser than 0) and no-interaction (if the interaction values was equal to 0). Those values were consequently replaced by 1, -1 and 0 respectively using the *ι*(*θ*) function defined in equation 2.

The Ancestors Variance Index (AVI) only considers interactions between a given gene and its parents (i.e. the genes regulating that given gene), while the Descendants Variance Index (DVI) only considers interactions with its children (i.e. genes regulated by that gene). Those two indices make it easy to identify if a gene’s interactions vary mostly because of the interactions with genes upstream or downstream.

In particular, a gene KO is expected to have an effect on downstream genes.

For each gene *i* in a collection of GRNs, the indices were defined as :

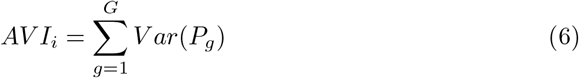

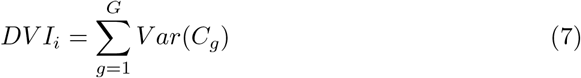

where *P*_*g*_ is the vectors of interaction values between gene *g* and its parents and *C*_*g*_ the vector of interactions values with its children. *G* is the number of genes in the GRNs.

### Measure of distance between multivariate distributions

Distances between multivariate distributions (simulated or experimental data) were measured using the Kantorovich distance [10] (also referred to as Wassertein or EMD distance). Because of the high number of variables (i.e. genes), the number of sample points (i.e. cells) was however too low to correctly estimate the multivariate Kantorovich distance. We thus devised a modified d istance w hich c omputes t he s um of Kantorovich distances on marginals (i.e. one variable at the time), making it practically usable.

This distance, named *Kantorovich*_1*D*_, has the following form :

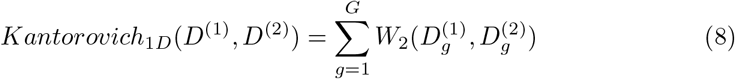

where *D*^(1)^ and *D*^(2)^ are 2 multivariate distributions (both with the same G vari-ables) and *W*_2_ is the regular Kantorovich distance. 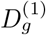 and 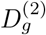 refer to the vector of values in *D*^(1)^ and *D*^(2)^ for variable *g*.

### Simulation balancing

Before any simulation could be executed, it was essential to correctly balance it, i.e. constant hyper-parameters had to be chosen such that the initial state produced by the simulation was a desired stable state. Here, parameters *β*_*Kon*_ and *β*_*Koff*_ act as adjustment variables which can force genes into high or low expression regimes by increasing or decreasing the value of the interaction function *ϕ*.

Finding the correct *β* values is a non trivial task since 2 parameters (*β*_*Kon*_ and *β*_*Koff*_) need to be fitted for each gene of the *G* genes in the GRN. In our case, 49 *×* 2 = 98 parameters needed to be fitted per GRN. Such high dimensional optimization problems suffer from the curse of dimensionality and either converge to a solution in very long times or don’t allow a solution to be found at all.

Fortunately, each gene could be considered independent from the other since the goal of the balancing process was to find *β* values such that the expression level of all genes remained totally unchanged. In this case, all mRNA and protein distributions (apart from that of the gene we are trying to balance) can be considered constant through time, thus transforming a *G ×* 2 dimensional optimization problem into *G* 2 dimensional problems.

Resolution of these problems was however made difficult by the stochastic nature of the simulation outputs. To that end, we adapted a simulated annealing algorithm to the noisy cost function case as described in [21] and [22]. We designed a cost function taking as input a tuple of *β*_*Kon*_ and *β*_*Koff*_ values, which executes the simulation of a single gene with those *β* values for 20 hours and returns the Kantorovich distance between the simulated data (at t=20h) and the initial data (at t=0h). The simulated annealing method is detailed in Additional file 1.

### Simulation initialization

After balancing, simulations were initialized by setting mRNA counts for each gene from the distribution of single-cell RT-qPCR data which was used in the GRN inference task. Only data at the initial time point was used here.

### Measure of information gained after perturbation

Simulations of perturbations on the set of candidate GRNs predicted effects on varying numbers of genes and significant gene level variations were observed for different proportions of the 364 GRNs. To determine which perturbation carried the most information, we computed the entropy of the proportion of GRNs with significant expression variation for each of the 49 genes. As an example, *FNIP1* had significant expression variation for genes *SNX22* (in 88 out of the 364 GRNs), *SLC25A37* (in 45 GRNs) and *ENSGALG00010025565* (in 102 GRNs). We thus computed the entropy using equation 9, where *p* was the vector 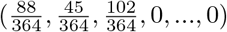 (with 46 trailing zeros for the 46 genes with no significant variation) encoding the proportion of GRNs with expression variation.

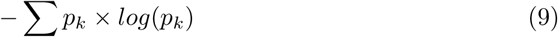

### Cell culture

T2EC were extracted from 19-days-old SPAFAS white leghorn chicken’s embryos’ bone marrow (INRA, Tours, France). These primary cells were maintained in self-renewal in LM1 medium as previously described [9].

### CRISPR plasmids construction

A guide RNA against FNIP1’s sequence (ENSGALG00000017462) was designed using the CRISPOR design tool [23] to target the exon number 5. Oligonucleotides were purchased from Eurogentec (Table 3). The guide was cloned after hU6 promoter into BbsI-digested pCRISPR-P2A-tRNA vector [24]. The efficiency of our CRISPR vector in T2EC cells was confirmed by analyzing mutations after sequencing.

**Table 3.**
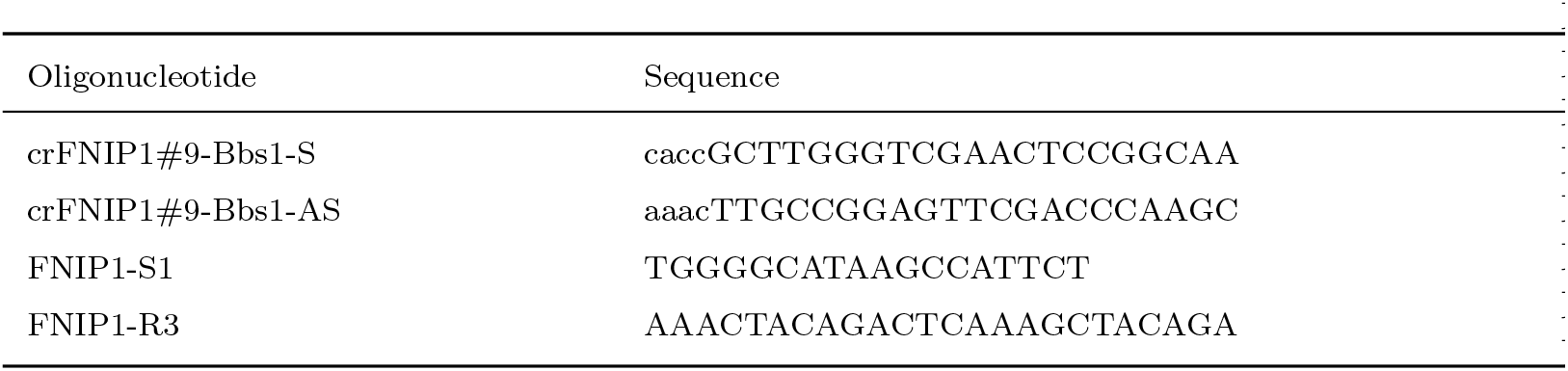
Oligonucleotides sequences used for CRISPR plasmid construction.

### Cells transfection

After 12 days in culture, 30 *×* 10^5^ cells were resuspended in 100µL of transfection medium (Cell Line Nucleofector Kit V – Amaxa) and transfected with 6,5µg of pCRISPR-P2A-tRNA empty vector or pcrFNIP#9 vector using the T16 K652 program. 500µL of RPMI (RPMI 1640 Medium no phenol red - Gibco) were added to the cell solution for a recovery step of 8 min. Then cells were transferred at 1, 25.10^6^ cell-s/ml in LM1 medium without penicillin and streptomycin and grown in standard culture conditions.

### Single cells sorting

24H after transfection, cells were harvested and resuspended in LM1 medium. Sorting was performed at room temperature using BD FACS Aria 1 flow cytometer. Living GFP-expressing cells were sorted in 96 wells U-shape culture plates containing 50µL of regular LM1. Non-transfected cells were also sorted to be used as a negative control. Plates were then placed back in incubator at 37°C, with 5% CO2 (Fig 3).

### Identification of *FNIP1* KO clones by sequencing

30 clones were selected 7 days post-sorting and half of the culture was collected for DNA/RNA extraction with Quick-DNA/RNA™ Microprep Plus Kit (Zymo) according to the manufacturer’s protocol. The amplified DNA fragments (with FNIP1-S1/FNIP1-R3 primers (Table 3)) were cloned into the pCR™4-TOPO® TA vector (TOPO™ TA Cloning™ Kit for Sequencing without competent cells - Invitrogen). We selected a clone presenting a frame shift leading to an early stop codon on both alleles for subsequent transcriptomics analysis.

### Single-cell RT-qPCR analysis

Individual cells from clones transfected with the pcrFNIP#9 vector (*FNIP1* KO cells) or the empty vector clones (wild type cells) were sorted into 96-well plates using BD FACS Aria 1 flow cytometer. All the manipulations related to the high-throughput scRT-qPCR experiments in microfluidics were performed according to the protocol recommended by the Fluidigm company (PN 68000088 K1, p.157-172). All steps from single-cell isolation, gene selection, data generation by scRT-qPCR are described in details in [25].

### Declarations

#### Ethics approval and consent to participate

According to the directive 2010/63/eu of the European Parliament and of the Council of 22 September 2010 on the protection of animals used for scientific purposes, ‘procedure’ excludes the killing of animals solely for the use of their organs or tissues. Since this is exactly what has been done in the present paper, since no procedure was involved, no approval was required.

## Consent for publication

Not applicable

## Availability of data and materials

The datasets analysed during the current study are available in the OSF repository, https://osf.io/r3ujs/. Additionally, TopoDoE is made available as a Python library which can be downloaded from github at https://github.com/Vidium/topodoe. The simulated annealing algorithm modified for noisy cost functions that we used for balancing our simulations is also available as a Python library. Source code can be downloaded from github at https://github.com/Vidium/josiann. The library can also be installed with pip by running ‘pip install josiann’.

## Authors’ contributions

MB designed and developed TopoDoE and Josiann. SZ, EV, CF, OG and SG designed the experimental process and generated the data analyzed in this work. MB, AB and OG drafted the manuscript. All authors read and approved the final manuscript.

## Acknowledgments

We would like to thank all members of the SBDM, Dracula and Vidium teams for their valuable advice and fruitful discussions. We also thank Alice Hugues, Emma Risson and Kaushik Karambelkar for participating as interns to the very early phases of that work. We thank the BioSyL Federation and the LabEx Ecofect (ANR-11-LABX-0048) of the University of Lyon for inspiring scientific events.

## Supplementary Information

### Additional_file_1.pdf

Title: Supplementary methods. Description: A description of Josiann, a noisy simulated annealing algorithm implementation in Python.

### Additional_file_2.pdf

Title: Supplementary material. Description: supplementary figures and tables.

