## Additional file 1 for "TopoDoE: A Design of Experiment strategy for selection and refinement in ensembles of executable Gene Regulatory Networks"

### Supplementary methods: Josiann, a simulated annealing algorithm for minimizing functions under noise

#### General description

Finding the minimum of a function is a classical optimization problem. However, when the data gathered for an optimization problem is noisy – because of experimental noise on the measurement or because of an underlying stochastic process, as it is the case with our simulations – or when an exhaustive evaluation of the cost function is too expensive, simple optimization methods might fail. Here we describe a modified simulated annealing procedure, adapted from [1], which takes random samples of the data to compute approximations of the cost function to minimize. The number of samples progressively increases to guaranty convergence. Because our cost function was expensive to compute, we implemented a series of computational optimizations to reduce the number of function evaluations as much as possible. The procedure was called Josiann (for Just anOther SImulated ANNealing) and made freely available at https://github.com/Vidium/josiann in the form of a Python package.

In the original definition of Simulated Annealing (SA) [2], an iterative procedure is applied to converge in probability to the global solution to the general problem :

$$\underset{x \in S}{\text{minimize}} \quad f(x) \tag{1}$$

where $f : S \to \mathbb{R}$ and $S \subset \mathbb{R}^d$ is a finite search space, i.e. a set of values at which $f$ can be evaluated. The number of dimensions $d$ can be large, but high dimension spaces make the convergence more difficult.

**Algorithm 1** Simulated Annealing algorithm

$x \leftarrow x_0$
$c \leftarrow f(x)$
$T \leftarrow T_0$
$k \leftarrow 0$

**while** $k \leq K$ **do**
  generate candidate position : $x_k \leftarrow Move(x)$
  $c_k \leftarrow f(x_k)$

  **if** $exp\left(\frac{c-c_k}{T}\right) > uniform(0,1)$ **then**
    accept candidate position : $x \leftarrow x_k$
    $c \leftarrow c_k$
  **end if**

  update temperature : $T \leftarrow update\_temperature(T)$
  $k \leftarrow k+1$

**end while**

The SA algorithm is described in algorithm 1. Briefly, it is initialized with a random position $x$, an associated cost $c = f(x)$ and an initial Temperature value $T^0$. At each iteration $k$, a candidate position $xk$ is generated by a selected 'move function', and the cost of that position $ck$ is computed by evaluating $f(xk)$. Then, the acceptance function is computed to determine if the candidate position is accepted or rejected. The acceptance function is dependant on the Temperature $T$ value and has the following characteristics :

- the candidate position is always accepted if $c^k < c$
- the candidate position is also accepted with probability $exp(\frac{c-c^k}{T^k})$ ($uniform(0,1)$ generates a value between 0 and 1 from a uniform distribution.)

Finally, $x$ and $c$ are updated if the candidate position is accepted and $T$ is decreased according to a cooling schedule (such as $T = T^0\alpha^k, 0.8 \leq \alpha \leq 0.9$).

It must be noted that for initial, large values of the Temperature $T$, a large fraction of candidate positions will be accepted, allowing the procedure to explore a good portion of the search space. As T decreases, positions with decreasing costs will be selected more and more, to allow the algorithm to converge towards a solution.

#### Balancing problem

In this study, before simulating any GRN, we had to calibrate one $\beta_{Kon}$ and $\beta_{Koff}$ parameter per gene. We defined a cost function as a function $f(x)$ in $\mathbb{R}^2$ where $x$ was the vector $(\beta_{Kon}, \beta_{Koff})$ which returned a real value. For each call to the function, a distribution of mRNA values also had to be supplied so that initial values could be set for all genes in the GRN. Then $\beta_{Kon}$ and $\beta_{Koff}$ of the gene of interest were set from vector $x$ ; expression data was obtained from our executable model simulated for 20 hours while forcing expression values for all genes besides the one of interest to remain constant ; and finally the Kantorovich distance of initial and simulated data at t=20h was computed and returned. The central role of the stochastic simulation of gene expression data meant that the cost function was noisy. The challenge of optimizing such functions was addressed by the algorithm described in the following section.

#### Convergence for noisy cost functions

When the cost function to evaluate is stochastic (i.e. we do not measure $f(x)$ directly but rather $\tilde{f}(x) = f(x) + \epsilon_x$, where $\epsilon_x$ is a value drawn from a random variable), the convergence of the regular SA algorithm is no longer guarantied. [1] proposed a modified version of that algorithm which also converges in the noisy setting. The core idea is that $f(x)$ can be approached with sufficient precision by computing the average of several evaluations of $f(x)$ :

$$\widehat{f}(x) = \frac{1}{n}\sum^{n} f(x) \quad (2)$$

##### Conditions of convergence

The modified SA algorithm effectively converges under some conditions, as demonstrated in [1] :

1. the noise $\epsilon_x$ must be generated by a symmetric distribution centered around zero.
2. the standard deviation $\sigma^k$ of the noise must decrease on the order of $O(k^{-\gamma})$ (with $k$ the number of iterations and $\gamma > 1$ an arbitrary constant)

As proved in [3], such conditions are met when $\sigma^k < T^k$ which leads to a sufficient general form of $\sigma^k$ :

$$\sigma^k = T^k(1-\epsilon)^k, \text{ with } 0 < \epsilon < 1 \tag{3}$$

with the general cooling schedule :

$$T^k = T^0\alpha^k, \text{ with } 0 \leq \alpha \leq 1 \tag{4}$$

Above results allow to derive equation 5 for computing the minimal number $n^k$ of required function evaluations at iteration $k$

$$n^k = \frac{N\sigma_{max}^2}{(N-1)\sigma^{2^k} + \sigma_{max}^2} \tag{5}$$

where $N$ is the maximum number of function evaluations and $\sigma_{max}^2$ is the worst-case value of $\sigma^2$ (i.e. the value of $\sigma^2$ at $k = 0$). As it is difficult to compute the variance of the Kantorovich distance, we tested multiple values of N and found that $N \geq 10$ gave satisfying results.

#### Move function

At each iteration, a new candidate position must be generated by a function which we call 'move function'. Many such functions have be proposed such as the original Metropolis (first described in [4] and used in the context of SA by [2]) or more complex ensemble moves where multiple points are used together to compute new candidates (for example see [5]). Sometimes, it is beneficial to work with discrete move functions. In this work, evaluations of the cost function were expensive but precise identification of the unknown parameters $x$ was not required as close values produced very similar outcomes. We thus implemented a move function which uses a set of allowed values $A = a_1, a_2, ..., a_p$ to compute a new position for each component of $x$. To do so, the candidate position for each element $x_i$ of $x$ is drawn from neighboring values in $A$, i.e. when $x_i$ is equal to $a_k$ in $A$, a new position is drawn with equal probability in $a_{k-1}, a_{k+1}$. When $x_i$ hits a boundary $a_1$ or $a_p$, the new position is chosen with probability 1 to be $a_2$ or $a_{p-1}$ respectively. This choice of move function greatly reduces the size of the search space, especially for optimization problems with few dimensions, which allowed to further reduce the required computation time, as described in the next section.

#### Other optimizations

Because the cost function to optimize required very expensive simulations to be carried out, we implemented optimization which allowed to reduce as much as possible the number of function evaluations that would be needed.

##### Caching of function evaluations

Because we used a move function producing discrete values and the number of dimensions were low, there was a high probability for the SA algorithm to visit the same position $x$ multiple times. Since function evaluations were assumed to be randomly distributed around the true cost, each evaluation was stored in a cache. In that way, only $n^k - l$ evaluations were required instead of $n^k$, where $l$ is the number of cached function evaluations at the same position computed in iterations 0 to $k-1$.

##### Detection of convergence

As the Temperature parameter drops, the fraction of accepted candidate moves drops to select positions with lower costs. When a clear minimum was found, we observed that a significant fraction of the last iterations were spent repeatedly computing the cost functions at neighboring positions while always being rejected. To detect such cases of early convergence, we compute the Root-Mean-Square Deviation (RMSD) of the costs of the last $w$ accepted positions, where $w$ is an hyper-parameter to be chosen and usually set to 50. When the RMSD dropped below 0.001, convergence was declared to have occurred and the SA algorithm was stopped early. This convergence detection allowed to reduce the number of iterations effectively computed by as much as one fifth in some cases.

##### Parallelization

Since $\beta$ values had to be calibrated for each gene in a GRN independently, a full GRN calibration in sequence was still a slow process. To address this issue, we implemented a parallel version of the algorithm in which the cost function now accepts a matrix $X_{2\times G}$ as input parameter, where each column $i$ in $X$ holds the $\beta_{Kon}$ and $\beta_{Koff}$ values for gene i. A modified simulation algorithm was used, in which all genes could be simulated independently in parallel. Finally, $G$ Kantorovich distances were computed. In this parallel mode, all genes are treated independently but can all be calibrated at once, greatly reducing the computation time needed.
