## Additional file 2 for "TopoDoE: A Design of Experiment strategy for selection and refinement in ensembles of executable Gene Regulatory Networks"

### Supplementary material

| short name | Ensembl gene ID | short name | Ensembl gene ID |
|---|---|---|---|
| AACS | ENSGALG00000002899 | ABCG2 | ENSGALG00000005792 |
| ACSL6 | ENSGALG00000006644 | ACSS1 | ENSGALG00000008632 |
| ANGPTL4 | ENSGALG00000000619 | ARHGEF2 | ENSGALG00000006840 |
| BATF | ENSGALG00000010323 | BCL11A | ENSGALG00000007866 |
| BPI | ENSGALG00000006756 | CD151 | ENSGALG00000006856 |
| CRIP2 | ENSGALG00000026471 | CTCF | ENSGALG00000001817 |
| CYP51A1 | ENSGALG00000009365 | DHCR7 | ENSGALG00000004106 |
| DHCR24 | ENSGALG00000010798 | DPP7 | ENSGALG00000009105 |
| EGFR | ENSGALG00000012363 | FAM208B | ENSGALG00000008171 |
| FHL3 | ENSGALG00000001516 | FNIP1 | ENSGALG00000017462 |
| GAB1 | ENSGALG00000009898 | GSN | ENSGALG00000001446 |
| HMGCS1 | ENSGALG00000014862 | HRAS1 | ENSGALG00000006885 |
| HSD17B7 | ENSGALG00000002681 | HSP90AA1 | ENSGALG00000011351 |
| LCP1 | ENSGALG00000016986 | LDHA | ENSGALG00000006300 |
| MFSD2B | ENSGALG00000016498 | MID2 | ENSGALG00000003860 |
| MVD | ENSGALG00000007267 | NSDHL | ENSGALG00000007493 |
| PIK3CG | ENSGALG00000008081 | PLS1 | ENSGALG00000002647 |
| REXO2 | ENSGALG00000007024 | RSFR | ENSGALG00000027165 |
| RUNX2 | ENSGALG00000026484 | sca2 | ENSGALG00000016152 |
| SCD | ENSGALG00000005739 | SLC6A9 | ENSGALG00000010098 |
| SLC25A37 | ENSGALG00000000378 | SNX22 | ENSGALG00000002206 |
| SQLE | ENSGALG00000016331 | STARD4 | ENSGALG00000000241 |
| SULF2 | ENSGALG00000004593 | TBC1D7 | ENSGALG00000012731 |
| TNFRSF21 | ENSGALG00000016719 | ENSGALG00010025565 | ENSGALG00010025565 |
| VDAC3 | ENSGALG00000003755 | | |

**S1 Table List of gene names and corresponding Ensembl gene IDs in the 364 candidate GRNs.**

| Test hypothesis | Gene name | p-value | q-value |
|---|---|---|---|
| | STARD4 | 0.9931 | 0.9996 |
| | SLC25A37 | 0.0000 | 0.0000* |
| | ANGPTL4 | 0.9685 | 0.9996 |
| | GSN | 0.9700 | 0.9996 |
| | FHL3 | 0.9961 | 0.9996 |
| | CTCF | 0.5773 | 0.9996 |
| | SNX22 | 0.0001 | 0.0011* |
| | PLS1 | 0.3583 | 0.9996 |
| | HSD17B7 | 0.9973 | 0.9996 |
| | AACS | 0.7651 | 0.9996 |
| | VDAC3 | 0.3665 | 0.9996 |
| less | MID2 | 0.9924 | 0.9996 |
| | DHCR7 | 0.9900 | 0.9996 |
| | SULF2 | 0.9561 | 0.9996 |
| | SCD | 0.9961 | 0.9996 |
| | ACSL6 | 0.0741 | 0.5558 |
| | BPI | 0.9984 | 0.9996 |
| | ARHGEF2 | 0.9923 | 0.9996 |
| | CD151 | 0.9971 | 0.9996 |
| | HRAS1 | 0.8347 | 0.9996 |
| | REXO2 | 0.8648 | 0.9996 |
| | MVD | 0.9975 | 0.9996 |
| | NSDHL | 0.9514 | 0.9996 |
| | BCL11A | 0.8422 | 0.9996 |
| | PIK3CG | 0.9889 | 0.9996 |

| Test hypothesis | Gene name | p-value | q-value |
|---|---|---|---|
| | FAM208B | 0.7315 | 0.9996 |
| | ACSS1 | 0.9640 | 0.9996 |
| | DPP7 | 0.9849 | 0.9996 |
| | CYP51A1 | 0.9941 | 0.9996 |
| | SLC6A9 | 0.0001 | 0.0011* |
| | ENSGALG00010025565 | 0.0000 | 0.0000* |
| | BATF | 0.9496 | 0.9996 |
| | DHCR24 | 0.9656 | 0.9996 |
| | HSP90AA1 | 0.9419 | 0.9996 |
| | EGFR | 0.9956 | 0.9996 |
| | TBC1D7 | 0.0525 | 0.4725 |
| less | HMGCS1 | 0.9338 | 0.9996 |
| | sca2 | 0.9994 | 0.9996 |
| | SQLE | 0.9979 | 0.9996 |
| | MFSD2B | 0.8349 | 0.9996 |
| | TNFRSF21 | 0.8440 | 0.9996 |
| | LCP1 | 0.9712 | 0.9996 |
| | CRIP2 | 0.9996 | 0.9996 |
| | RUNX2 | 0.9850 | 0.9996 |
| | RSFR | 0.9120 | 0.9996 |
| | STARD4 | 0.0069 | 0.0248 |
| | SLC25A37 | 1.0000 | 1.0000 |
| | ANGPTL4 | 0.0315 | 0.0675 |
| | GSN | 0.0300 | 0.0675 |
| | FHL3 | 0.0039 | 0.0195 |

| Test hypothesis | Gene name | Gene Ensembl ID | p-value | q-value |
|---|---|---|---|---|
| | CTCF | 0.4227 | 0.5141 | |
| | SNX22 | 0.9999 | 1.0000 | |
| | PLS1 | 0.6417 | 0.7404 | |
| | HSD17B7 | 0.0027 | 0.0186 | |
| | AACS | 0.2349 | 0.3020 | |
| | VDAC3 | 0.6335 | 0.7404 | |
| | MID2 | 0.0076 | 0.0248 | |
| | DHCR7 | 0.0100 | 0.0300 | |
| | SULF2 | 0.0439 | 0.0823 | |
| | SCD | 0.0039 | 0.0195 | |
| | ACSL6 | 0.9259 | 1.0000 | |
| greater | BPI | 0.0016 | 0.0186 | |
| | ARHGEF2 | 0.0077 | 0.0248 | |
| | CD151 | 0.0029 | 0.0186 | |
| | HRAS1 | 0.1653 | 0.2188 | |
| | REXO2 | 0.1352 | 0.2028 | |
| | MVD | 0.0025 | 0.0186 | |
| | NSDHL | 0.0486 | 0.0872 | |
| | BCL11A | 0.1578 | 0.2188 | |
| | PIK3CG | 0.0111 | 0.0312 | |
| | FAM208B | 0.2685 | 0.3356 | |
| | ACSS1 | 0.0360 | 0.0704 | |
| | DPP7 | 0.0151 | 0.0378 | |
| | CYP51A1 | 0.0059 | 0.0241 | |
| | SLC6A9 | 0.9999 | 1.0000 | |

| Test hypothesis | Gene name | p-value | q-value |
| --- | --- | --- | --- |
| greater | ENSGALG00010025565 | 1.0000 | 1.0000 |
| | BATF | 0.0504 | 0.0872 |
| | DHCR24 | 0.0344 | 0.0704 |
| | HSP90AA1 | 0.0581 | 0.0968 |
| | EGFR | 0.0044 | 0.0198 |
| | TBC1D7 | 0.9475 | 1.0000 |
| | HMGCS1 | 0.0662 | 0.1064 |
| | sca2 | 0.0006 | 0.0135 |
| | SQLE | 0.0021 | 0.0186 |
| | MFSD2B | 0.1651 | 0.2188 |
| | TNFRSF21 | 0.1560 | 0.2188 |
| | LCP1 | 0.0288 | 0.0675 |
| | CRIP2 | 0.0004 | 0.0135 |
| | RUNX2 | 0.0150 | 0.0378 |
| | RSFR | 0.0880 | 0.1366 |

**S2 Table**: **Differential expression of T2ECs after FNIP1's Knock-Out** Gene expression levels of the KO cells compared to the wild type cells using one-sided T-tests (the 'less' hypothesis means that genes are down-regulated in the KO condition). The obtained p-values are reported in column 'p-value' and corrected p-values for multiple testing are reported in column 'q-value'. Asterisks in the table indicate significant q-values (q-value $< 0.01$).

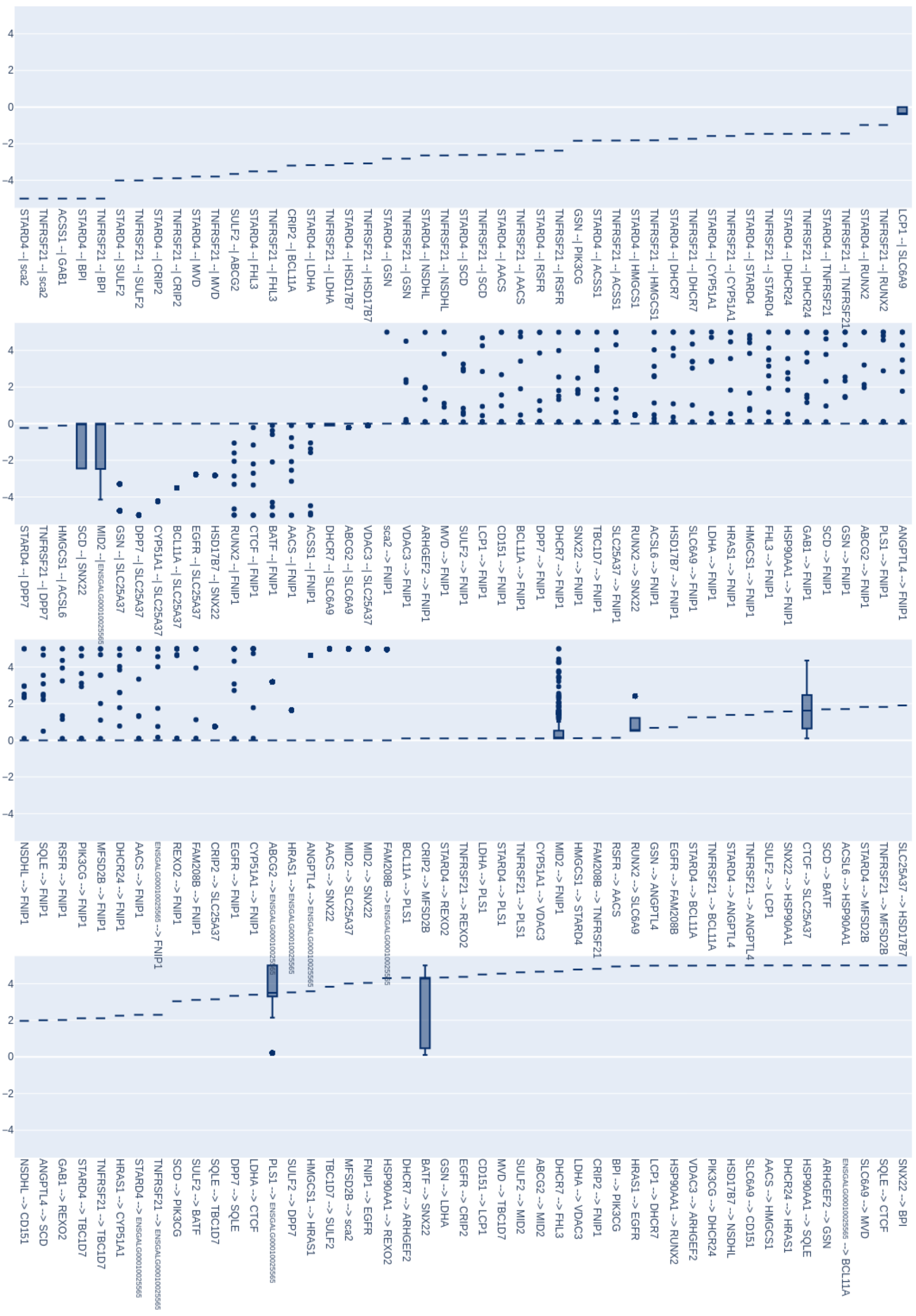




**S1 Fig Boxplots of interaction values.** Boxplots showing the distribution of interaction values for each stimulus-to-gene or gene-to-gene interaction which was non null in at least one of the 364 GRNs. Most interactions were identical (or close to being) in all GRNs, resulting in flat boxplots with few outliers.

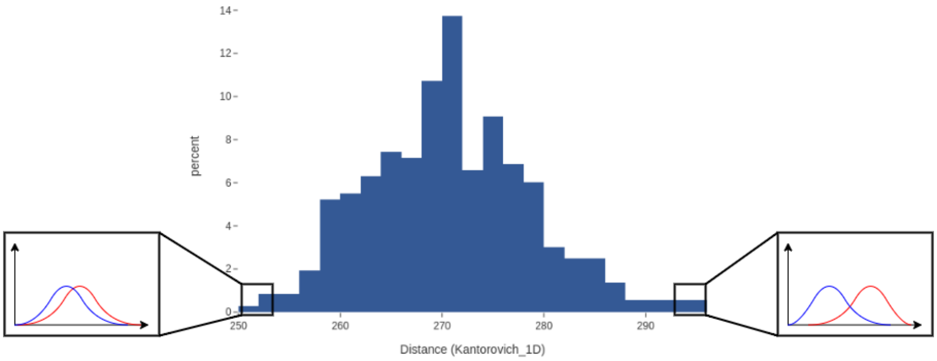


**S2 Fig Distance to reference data.** All 364 GRNs were simulated with the differentiation stimulus active for 72 hours and for each, mRNA expression data was recorded for 200 cells at time points 0h, 8h, 24h, 33h, 48h and 72h. Distributions of simulated expression data were compared to the reference experimental data (used in the inference process) using the $Kantorovich_{1D}$ distance. The histogram shows the distributions of the 364 obtained distances, where the leftmost bins hold the lowest distances (i.e. the distances of most similar distributions).

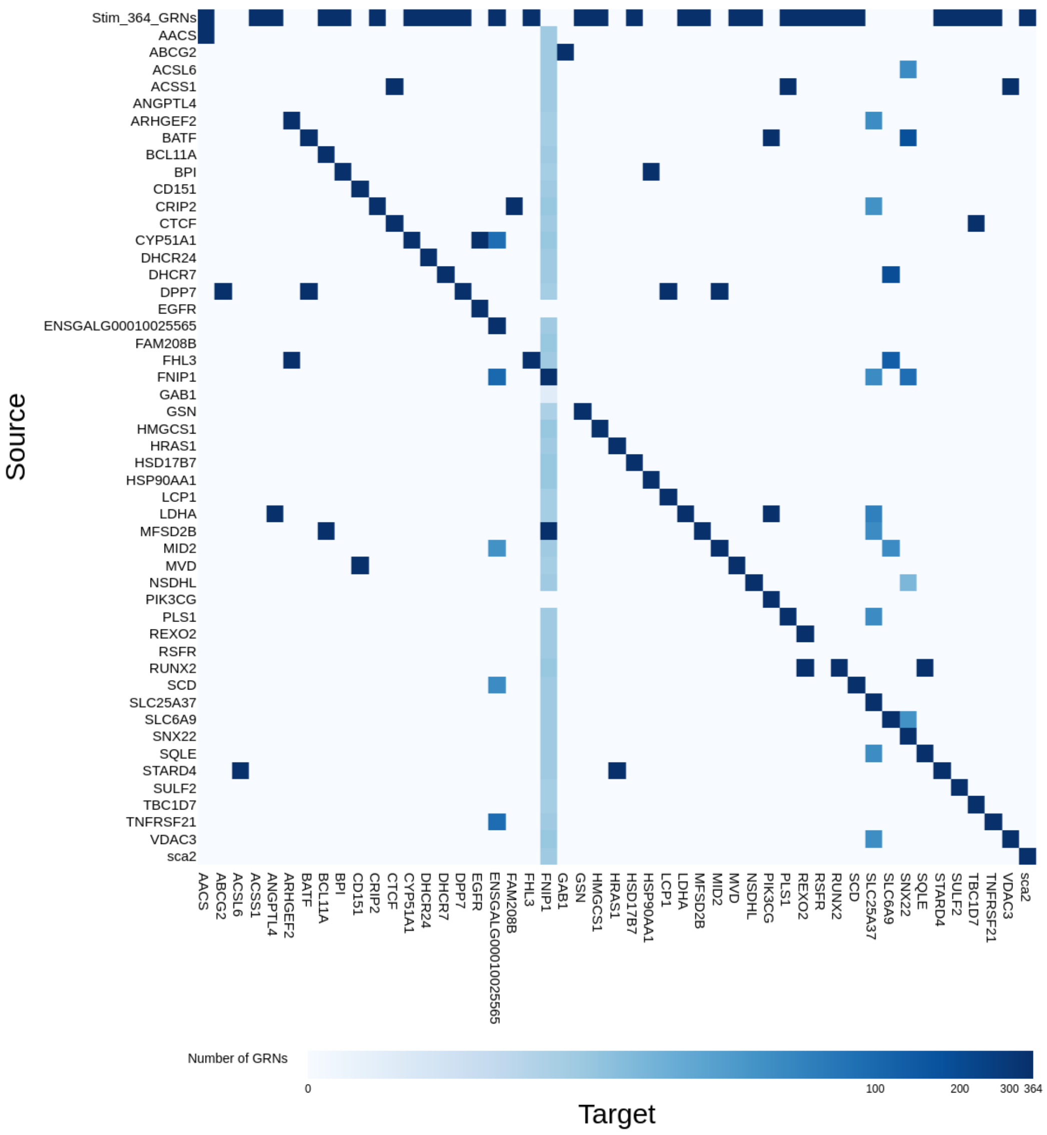


**S3 Fig Matrix of the interaction consensus.** Cells in the matrix indicate how many GRNs among the 364 candidates have an interaction between a gene (or stimulus) source (rows) and a gene target (columns). Values are color-coded on a log scale, where white indicates 0 and dark blue means 364.

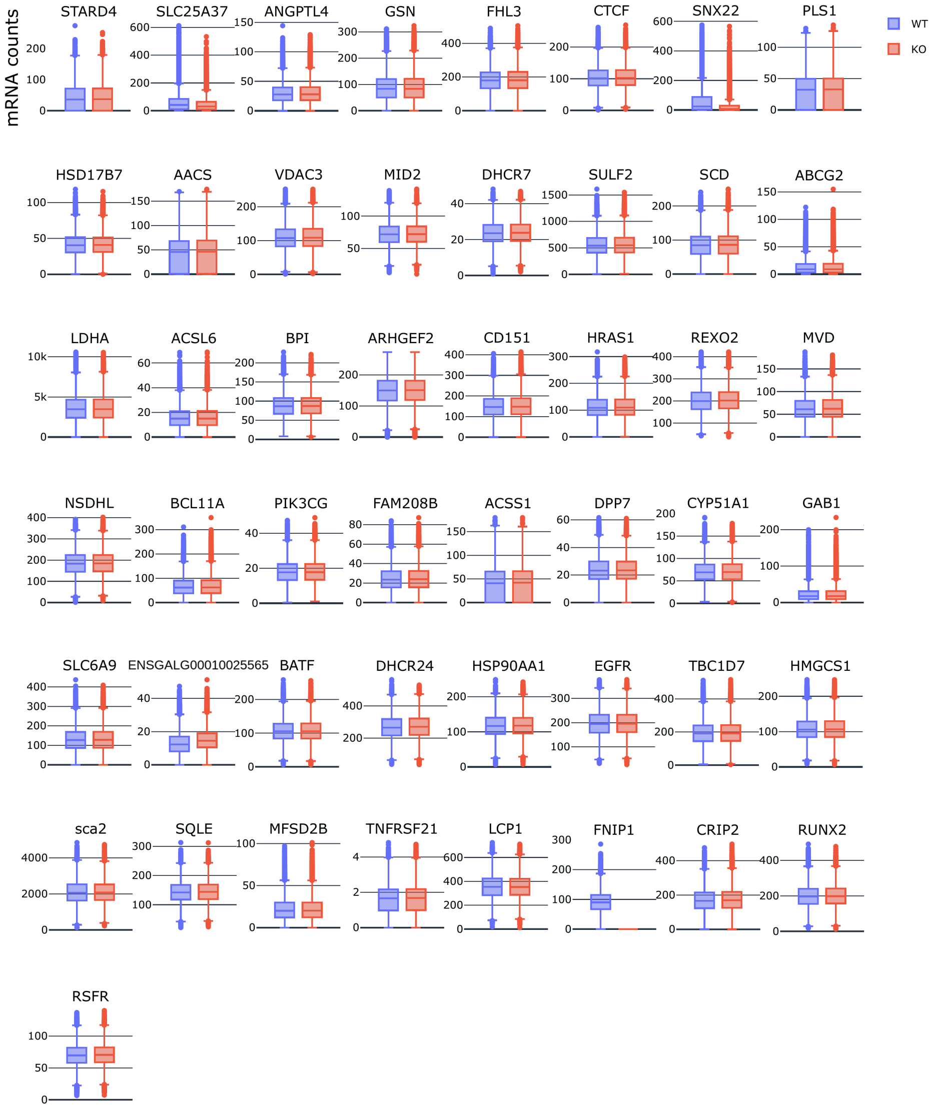


**S4 Fig Simulations of *FNIP1*'s Knock-Out.** Predicted expression variation for all 49 genes after *FNIP1*'s Knock-Out in all 364 GRNs. Wild type expression (no KO) is shown in **blue** while the KO condition is shown in **red**. Each box plot summarizes the expression level of 2000 simulated cells per each of the GRNs (728'000 data points per box plot in total).

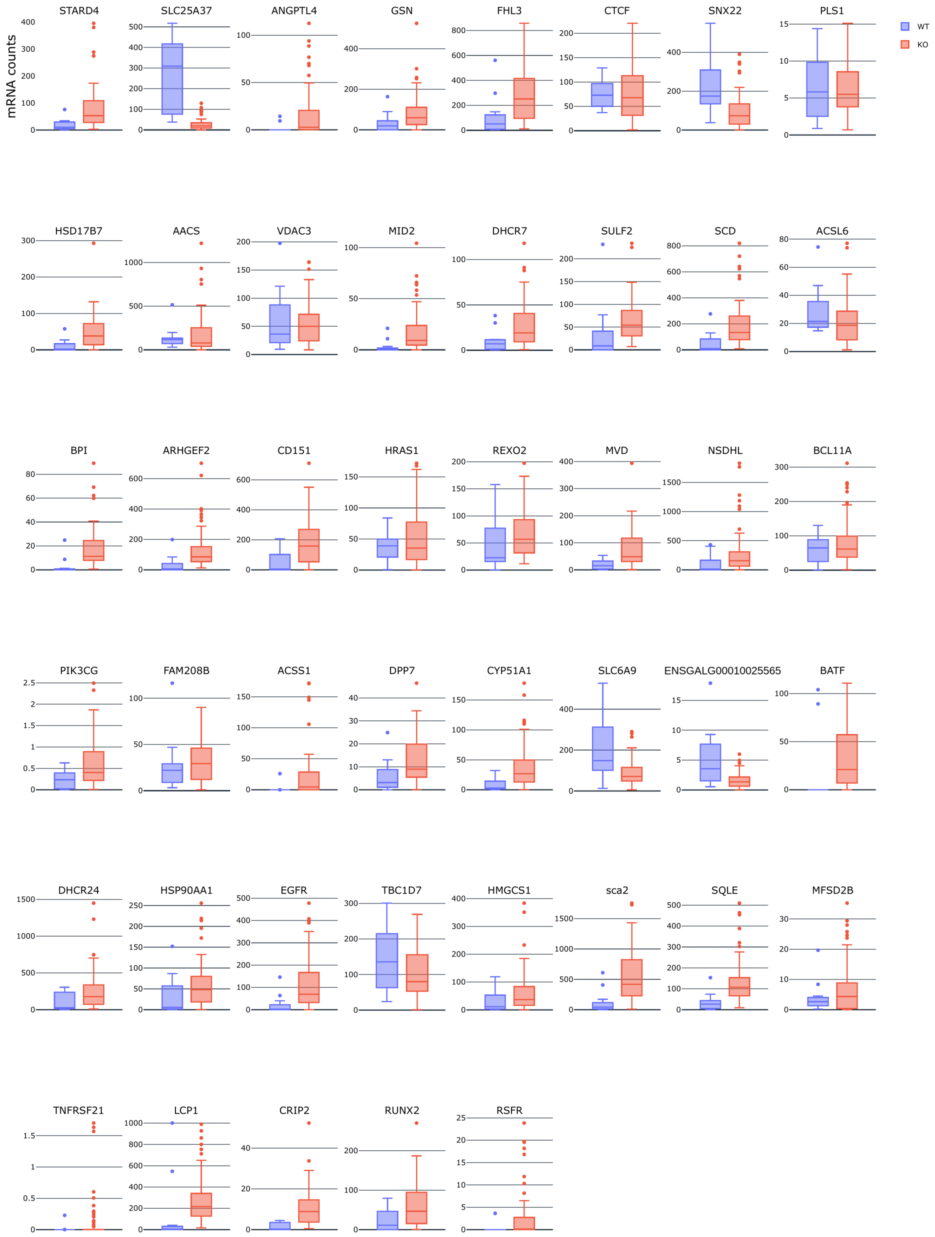


**S5 Fig Experimental data of T2ECs after *FNIP1*'s Knock-Out.** Observed expression variation for 45 genes (out of the 49 genes in the GRNs) after *FNIP1*'s Knock-Out in T2ECs. Missing genes are *ABCG2*, *LDHA*, *GAB1* and *FNIP1*. Wild type expression (no KO) is shown in **blue** while the KO condition is shown in **red**.

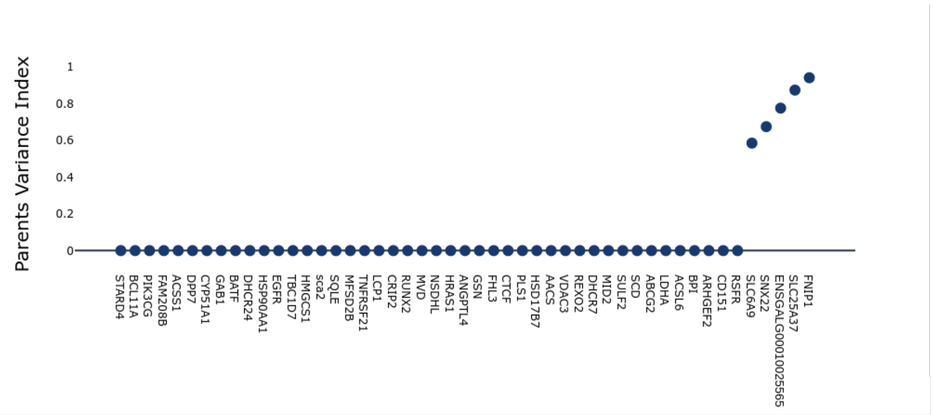


**S6 Fig  Ancestors Variance Index.** The index of ancestors variance is computed as the sum of the variance of the interaction values, in all candidate GRNs, between a gene and its regulators. A high value indicates that a gene has interactions that vary a lot among all the candidate GRNs.
